# Exogenous Sex Hormone Effects on Brain Microstructure in Women: A diffusion MRI Study in the UK Biobank

**DOI:** 10.1101/2020.09.18.304154

**Authors:** Leila Nabulsi, Katherine E. Lawrence, Vigneshwaran Santhalingam, Zvart Abaryan, Christina P. Boyle, Julio E. Villalon-Reina, Talia M. Nir, Iyad Ba Gari, Alyssa H. Zhu, Elizabeth Haddad, Alexandra M. Muir, Neda Jahanshad, Paul M. Thompson

## Abstract

Changes in estrogen levels in women have been associated with increased risk for age-related neurodegenerative diseases, including Alzheimer’s disease, but the impact of exogenous estrogen exposure on the brain is poorly understood. Oral contraceptives (OC) and hormone therapy (HT) and are both common sources of exogenous estrogen for women in reproductive and post-menopausal years, respectively. Here we examined the association of exogenous sex hormone exposure with the brain’s white matter (WM) aging trajectories in postmenopausal women using and not using OC and HT (HT users: n=3,033, non-users n=5,093; OC users: n=6,964; non-users n=1,156), while also investigating multiple dMRI models. Cross-sectional brain dMRI data was analyzed from the UK Biobank using conventional diffusion tensor imaging (DTI), the tensor distribution function (TDF), and neurite orientation dispersion and density imaging (NODDI). Mean skeletonized diffusivity measures were extracted across the whole brain, and fractional polynomial regressions were used to characterize age-related trajectories for WM microstructural measures. Advanced dMRI model NODDI revealed a steeper WM aging trajectory in HT users relative to non-users, and for those using unopposed estrogens relative to combined estrogens treatment. By contrast, no interaction was detected between OC status and age effects on the diffusivity measures we examined. Exogenous sex hormone exposure may negatively impact WM microstructure aging in postmenopausal women. We also present normative reference curves for white matter microarchitectural parameters in women, to help identify individuals with microstructural anomalies.

## INTRODUCTION

There are known sex differences in the risk for age-related neurodegenerative diseases such as Alzheimer’s disease (AD) (see Salminen et al., 2022 for review). On average, women over the age of 40 are at a greater risk of developing AD than age-matched men, and endogenous and exogenous estrogen levels also influence this risk in women (Shumaker et al. 2003; Savolainen-Peltonen et al. 2019; Song et al. 2020). Sex, as well as age, affects many aspects of grey and white matter (WM) microstructure across the lifespan (Cox et al. 2016; Nir et al. 2017; Ritchie et al. 2018; Toschi et al. 2020; Lawrence et al. 2021). Additionally, AD incidence and the menopause transition have frequently been linked, especially when the transition is induced surgically, or occurs at a younger age (Costantino et al. 2022)

Despite this evidence, women’s brain health is historically understudied (Covan 2005), and we know little about brain mechanisms underlying the reported sex differences in risk for degenerative diseases and how female-specific factors may influence women’s brain health throughout life. Given the growing population of older adults and the high percentage of females taking exogenous sex hormones worldwide, this is a major gap in the literature that requires immediate attention (Boyle et al. 2020).

Population based observational studies, as opposed to clinical trials, offer the large sample size, diversity, and statistical power required to investigate factors that influence aging trajectories across the lifespan. The UK Biobank is currently the largest prospective study of aging, gathering extensive questionnaires, physical and cognitive measures, neuroimaging data and biological samples in a population-based cohort of middle- to older-aged adults living in the United Kingdom (UK). Such large-scale population-based studies can identify subtle effects that could go undetected in smaller samples. Factors that modulate aging processes could inform future clinical trials and clinical decisions and guide recommendations for women at risk for age-related neurodegenerative disorders.

Some prior neuroimaging studies report relationships between levels of endogenous sex hormones - such as estradiol - and developmental processes, lifespan trajectories of aging (Salminen et al. 2022) and brain plasticity (Boyle et al., 2020; Galea et al., 2014; Simerly, 2002). Several brain structural changes have been associated with estrogen fluctuations in pregnant women (Hoekzema et al. 2017) and in pre-menopausal women across the menstrual cycle (Barth et al., 2016). Beneficial effects of endogenous estrogen levels have been reported generally for gray matter, with larger brain volumes for women during the reproductive years (den Heijer et al. 2003; Lisofsky et al. 2015). Even so, negative effects of endogenous estrogen levels on brain regional volumes have been reported in menopausal women (Resnick et al. 2003).

While evidence suggests that estrogen exerts a neuroprotective effect in reproductive and pre-menopausal women and unfavorable effects post-menopause, whether post-menopausal *exogenous* estrogen exposure is beneficial or detrimental for cognition is still an open question and warrants further investigation. Hormone therapy (HT) is a common source of exogenous estrogen in women menopausal years. A large randomized, double-blind, placebo-controlled clinical trial (N=4,894, Women’s Health Initiative Memory Study, WHIMS) of post-menopausal women (≥65 years) receiving estrogen plus progestin therapy reported for the first time that postmenopausal HT exposure may be detrimental for cognition (measured through 4 phases of neuropsychological and psychiatric assessments) and may increase their risk for probable dementia (Shumaker et al. 2003). A subsequent study using the WHIMS cohort reported greater brain atrophy in post-menopausal women treated with unopposed and conjugated estrogens (Resnick et al. 2003). Estrogen therapy is also called unopposed estrogen therapy because a second hormone (progestin) is not used along with the estrogen, whereas conjugated estrogen therapy includes a second hormone in addition to estrogen. A longitudinal study (Kantarci et al. 2016) reported increasing ventricular enlargement over 4 years in recently menopausal women (5-36 months past menopause) following conjugated estrogens. More recently, a large-scale UK Biobank population study of 16,854 middle to older-aged women (de Lange et al. 2020) reported that higher cumulative sex hormone exposure was associated with more adverse effects on the brain’s gray matter (i.e., more advanced *brain age*, a biomarker for aging); no information on HT treatment type was reported. In APOE⍰4 carriers, conjugated estrogens were found to reduce amyloid-β deposition (Kantarci et al. 2016), and higher levels of estradiol following HT were associated with less pronounced brain aging (de Lange et al. 2020); all relative to non-carriers. Other neuroimaging studies report a neuroprotective effect of unopposed estrogen treatment on WM (Ha et al. 2007).

The inconsistency of HT neuroimaging markers highlights the fact that there is still much to learn about the therapeutic potential of exogenous estrogen use. This variability and lack of consensus regarding HT effects on brain structure in women is a major gap in the literature. Large clinical trials suggest that the duration of HT and age of onset may be key factors in estrogen-mediated effects on the brain; the timing of the initiation of HT relative to menopause, age, or both, may relate to the health of the underlying vascular tissue and to other factors such as reduction in or down-regulation of estrogen receptors. Several clinical studies of HT exposure effects on the brain led to the critical period hypothesis, which states that HT may be neuroprotective if initiated near the time of cessation of ovarian function – that is, within around 5 years of menopause (MacLennan et al. 2006; Espeland et al. 2015). Consistent with the critical period hypothesis, in 65 women (63% receiving unopposed estrogen), Erickson and colleagues (2010) suggested that there is a limited window of opportunity for HT at the time of menopause, whereby shorter intervals between menopause and the initiation of treatment were associated with larger hippocampal volumes compared with longer intervals (Erickson et al. 2010). Other studies in favor of this hypothesis, such as KEEPS (a randomized double-blind placebo-controlled study; Gleason et al., 2015) indicated no adverse effects on measures of cognition in recently menopaused women, beneficial mood effects up to 4 years using conjugated estrogens, but not using estrogen only (oral 17β-estradiol). The ELITE study (randomized double-blind placebo-controlled study; Hodis et al., 2016) also indicated that women prescribed HT (oral 17β-estradiol) much later (>10 years after menopause) may be at higher risk of AD, as opposed to commencing therapy within 6 years after menopause. Findings by Gleason et al., (2015) and Hodis et al., (2016) were perhaps mediated by women’s cardiovascular risk profiles (Gleason et al. 2015; Hodis et al. 2016). In the UK Biobank genetic factors were also found to contribute to how timing of treatment initiation influenced women’s brain aging trajectories (de Lange et al. 2020); indicating beneficial effects of HT initiation before onset of menopause in APOE⍰4 carriers, relative to non-carriers. However, a study using the WHIMS data (conjugated estrogens; (Espeland et al. 2015) showed that conjugated therapy delivered near the time of menopause provided no detectable cognitive benefit or detriment.

Traditionally, research on steroid actions in the brain has focused mainly on postmenopausal HT and has rarely controlled for the use of hormonal oral contraception (OC). The effects of synthetic steroids contained OC on brain and cognitive abilities is understudied, despite the increasing use of OC globally (World Health Organization 2019). OC have been on the market for over 50 years and used by 100 million women (Christin-Maitre 2013; Pletzer and Kerschbaum 2014; Daniels and Abma 2019; World Health Organization 2019). Contemporary OC literature has focused mainly on comparing cognitive performance in users and non-users across the menstrual cycle of pre-menopausal women (Mordecai et al. 2008; Gogos 2013). Even so, these studies are inconclusive due to the small sample sizes considered. One study in postmenopausal women indicates that OC usage was not associated with more evident brain aging (de Lange et al. 2020).

There is a need for large-scale studies to expand on previous research and further evaluate sex-hormone factors specific to women in the context of aging on the brain. Here, for the first time, we set out to measure and model age-associated effects on WM microstructure to infer possible effects of exogenous sex hormones on the brains of postmenopausal women by using multiple diffusion MRI (dMRI) models.

Diffusion MRI is an *in vivo* imaging technique that allows quantitative investigation of microstructural differences in WM tracts by exploiting the Brownian motion of water molecules, and the dependency of this diffusion process on the cellular environment (Jones 2008). Late-life cognitive decline may be caused, in part, by microstructural deterioration in the brain’s neural pathways. Processes such as neuronal and glial cell loss, impaired myelin production, and axonal demyelination, may impair information transfer efficiency in the brain’s WM networks. The WM *disconnection theory* is largely supported by emerging data from dMRI studies of the brain, in both healthy and diseased subjects (Bennett and Madden 2014; Pievani et al. 2014; Lawrence et al. 2021). Thus, an accurate characterization of how and where the brain’s WM microstructure changes with age is a vital goal in brain aging research.

Here, we aimed to address the gap in variability and lack of consensus regarding exogenous sex hormone effects on brain structure in women by examining the association of HT and OC exposure with the brain’s white matter aging trajectories in postmenopausal women. We explored this association using different dMRI models and accounting for several clinical variables and confounding factors. Drawing on the existing literature on sex hormone effects on the brain, we hypothesized that sex hormone therapy would *negatively* impact women’s WM aging. We followed up with exploratory analyses to evaluate differences in HT regimen type (unopposed or conjugated estrogens), as these were hypothesized to differentially impact WM aging trajectories in women based on previous literature (Gleason et al. 2015; Savolainen-Peltonen et al. 2019). Further analyses were carried out to evaluate how WM aging relates to the duration of therapy, age at therapy onset and age at menopause, in women. We analyzed single- and multi-shell dMRI brain data from the UK Biobank (Miller et al. 2016a) and processed them using 3 increasingly complex dMRI models. We hypothesized that more sophisticated dMRI models that relate the signals from diffusion MRI to geometric models of tissue microstructure, as opposed to traditional tensor-based methods, will estimate more sensitively the microstructural complexity of aging WM trajectories in women using sex hormone therapy. Diffusion indices were extracted and averaged across a whole brain WM skeleton. We used fractional polynomial (FP) regression to characterize sex hormone effects on age-related trajectories for WM metrics. We also computed normalized centile curves to visualize WM aging trajectories for the major diffusivity metrics.

## MATERIALS & METHODS

### Sample

Our sample was drawn from the UK Biobank (Miller et al. 2016b) and included postmenopausal women aged 45-80 years old. We focused on the subsample of the overall UK Biobank that had neuroimaging data available for download at the time of writing. Exogenous sex hormone exposure in women was estimated from the HT and OC variables. Our primary analyses involved HT exposed women (N=3,033; ***Table 1***), where age effects on WM measures were compared to those in women who never took HT (N=5,093). Subsequently, we investigated age effects on WM measures in OC users (N=6,964, ***Table 2***), relative to those who never took OC (N=1,156). There was an overlap of N=748 women between HT and OC non-users; HT and OC models were run separately.

**Table 1.** Sample demographics for HRT users and non-users. Data shown as Mean ± standard deviation (SD), % for each variable in each of the groups, N = sample size; BMI = body mass index; HT = hormone replacement therapy; χ^2^-test = chi-squared test.

|  | HT users | non-users | t-test | p |
| --- | --- | --- | --- | --- |
| **Sample, N** | 3,033 | 5,093 |  |  |
| **Age at scan [years]**<br>*mean±SD*<br>*range* | 65.4±6.5<br>45.5-80.7 | 60.3±7.1<br>45.2-79.6 | 32.1 | <0.001 |
| **Age at Menopause**<br>*mean±SD* | 49.1±5.9 | 51.0±4.1 | 14.6 | <0.001 |
| **Number of live births**<br>*mean±SD*<br>*median*<br>*range* | 1.81±1.1<br>2<br>0-7 | 1.72±1.2<br>2<br>0-9 | 3.7 | <0.001 |
| **Waist-to-hip**<br>*mean±SD* | 1.6±0.5 | 0.81±0.5 | 3.4 | <0.001 |
| **Education [N]**<br>*high school / college* | 1290 / 1743 | 1713 / 3380 | 64.6 | <0.001 |
|  |  |  | **χ<sup>2</sup>-test** | **p** |
| **Menopause [N, %]**<br>*yes* | 2,998 (98.9%) | 4,508 (88.5%) | 250.5 | <0.001 |
| **Hysterectomy [N, %]**<br>*yes* | 404, 13.3% | 210, 4.1% | 677.7 | <0.001 |
| **Oophorectomy [N, %]**<br>*yes* | 535, 17.6% | 175, 3.4% | 491.4 | <0.001 |
| **Age at HT onset**<br>*mean±SD* | 47.8±5.6 | - | - | - |
| **Duration of HT therapy [years]**<br>*mean±SD*<br>*median*<br>*range* | 7.1±6.2<br>5<br>0.3-38 | - | - | - |
| **Diabetes**<br>*yes* | 113, 3.7% | 159, 5.2% | 101.7 | <0.001 |
| **Hypertension**<br>*Heart attack*<br>*Angina*<br>*Stroke*<br>*High blood pressure* | 15<br>29<br>20<br>447 | 17<br>24<br>19<br>515 | 48.9 | <0.001 |
| **Diagnosis of Major Depression Disorder**<br>*Probable recurrent major depression (severe)*<br>*Probable recurrent major depression (moderate)*<br>*Single probable major depression episode* | 73<br>160<br>92 | 93<br>237<br>116 | 101.1 | <0.001 |

**Table 2.** Sample demographics for OC users and non-users. Data shown as Mean ± standard deviation (SD), % for each variable in each of the groups, N = sample size; BMI = body mass index; OC = oral contraceptive; χ^2^-test = chi-squared test.

|  | OC users | non-users | t-test | p |
| --- | --- | --- | --- | --- |
| **Sample, N** | 6,964 | 1,156 | - | - |
| **Age at scan [years]**<br>*mean±SD*<br>*range* | 61.8±7.1<br>45.2-80.7 | 65.1±7.9<br>46.1-79.6 | 14.6 | <0.001 |
| **Age at Menopause**<br>*mean±SD* | 50.3±5.0 | 50.3±5.1 | 0.27 | 0.79 |
| **Number of live births**<br>*mean±SD*<br>*median*<br>*range* | 1.8±1.1<br>2<br>0-9 | 1.7±1.4<br>2<br>0-8 | 2.03 | 0.04 |
| **Waist-to-hip**<br>*mean±SD*<br>*median* | 0.81±0.7<br>0.80 | 0.81±0.07<br>0.81 | 0.26 | 0.797 |
| **Education [N]**<br>*high school / college* | 2,495 / 4,468 | 507 / 649 | 27.4 | <0.001 |
|  |  |  | **χ<sup>2</sup>-test** | **p** |
| **Menopause [N, %]**<br>*yes* | 5,327, 76.5% | 926, 80.1% | 4.1 | 0.04 |
| **Hysterectomy [N, %]**<br>*yes* | 496, 7.1% | 118, 10.2% | 13.4 | <0.001 |
| **Oophorectomy [N, %]**<br>*yes* | 599, 8.6% | 111, 9.6% | 1.2 | 0.23 |
| **Age at OC onset**<br>*mean±SD* | 20.8±4.3 | - | - | - |
| **Duration of OC therapy [years]**<br>*mean±SD*<br>*median*<br>*range* | 11.7±8.4<br>10<br>1-46.6 | - | - | - |
| **Diabetes**<br>*yes* | 232, 3.3% | 40, 3.5% | 101.4 | <0.001 |
| **Hypertension**<br>*Heart attack*<br>*Angina*<br>*Stroke*<br>*High blood pressure* | 28<br>48<br>32<br>791 | 4<br>5<br>7<br>172 | 48.9 | <0.001 |
| **Diagnosis of Major Depression Disorder**<br>*Probable recurrent major depression (severe)*<br>*Probable recurrent major depression (moderate)*<br>*Single probable major depression episode* | 148<br>350<br>182 | 18<br>47<br>27 | 86.0 | <0.001 |

We note that hormone exposure and hormone treatment are not randomized across individuals, as would be the case in a randomized clinical trial; instead, the study is a naturalistic epidemiological study of a large population (for assumptions and limitations of this approach, see *Discussion*).

### Analysis of Aging Trends

We modeled the effect of age on diffusion metrics (described in ***Table 3)*** in HT and OC users using higher order polynomial (FP) regressions (Royston and Altman 1994; Sauerbrei et al. 2006). Specifically, we used the multivariate fractional polynomial (*mfp*) package implemented in R (R Core Team, 2016). The *mfp* package used a predefined set of power terms (*-2, -1, -0*.*5, 0*.*5, 1, 2, 3*) and the natural logarithm function, and up to two power combinations to identify the best fitting model. FP for age was written as age^(p0.5, p1, …)^ where p in age refers to regular powers, and age^(o)^ refers to ln(age). Classic multiple regression models assume a linear relationship between the independent and dependent variables. However, age effects on the brain may be nonlinear; effects are the aggregate of a vast number of cellular processes, each with potentially different profiles of acceleration and decline. As the age effects are likely highly nonlinear, with a functional form that is not known *a priori*, fractional polynomials may offer a more adaptive modeling approach (Royston and Altman 1994). These have been used to chart age trajectories for volumetric and other structural measures by the ENIGMA Lifespan group (Dima et al. 2022; Frangou et al. 2022).

**Table 3.**
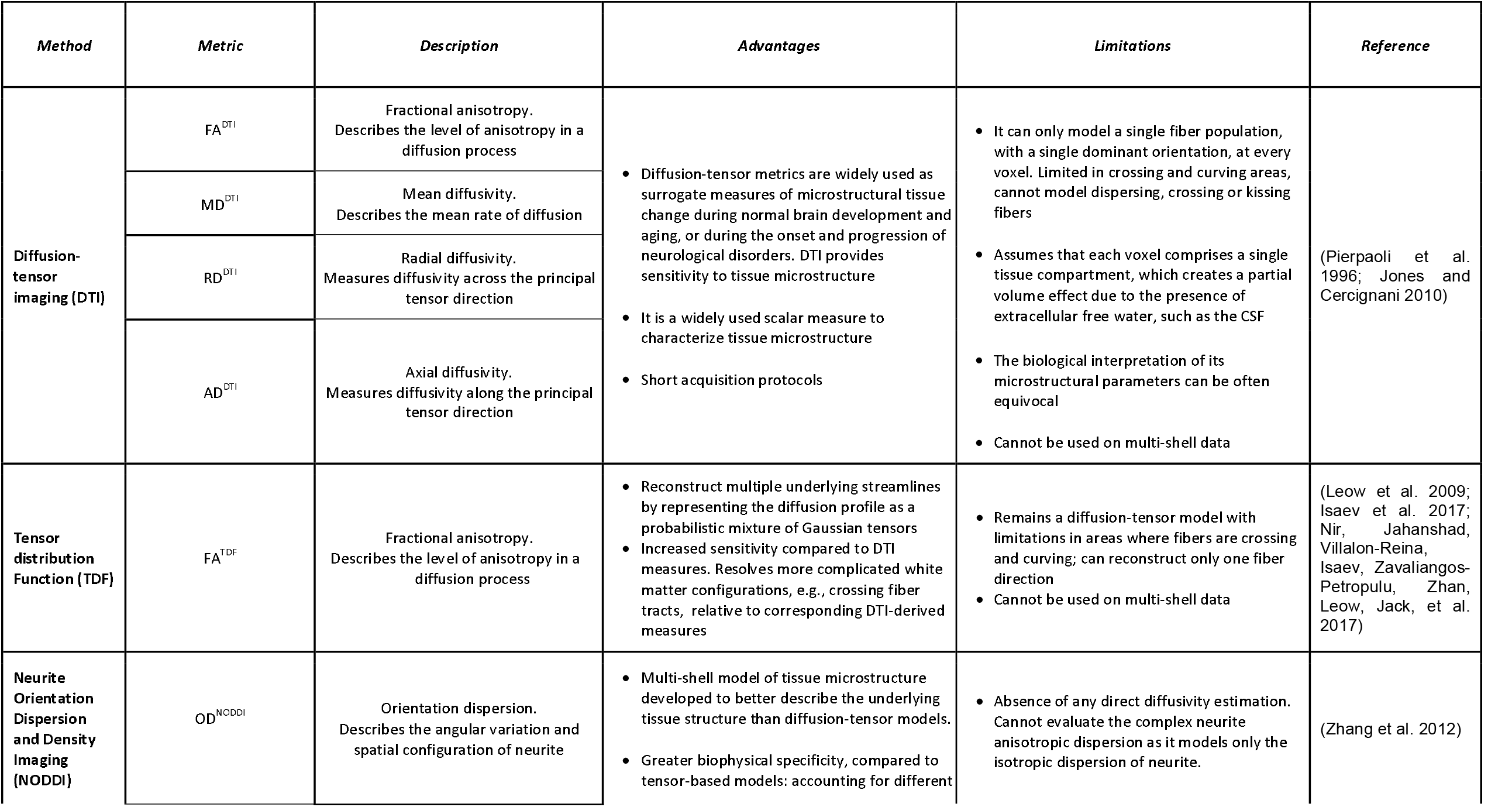
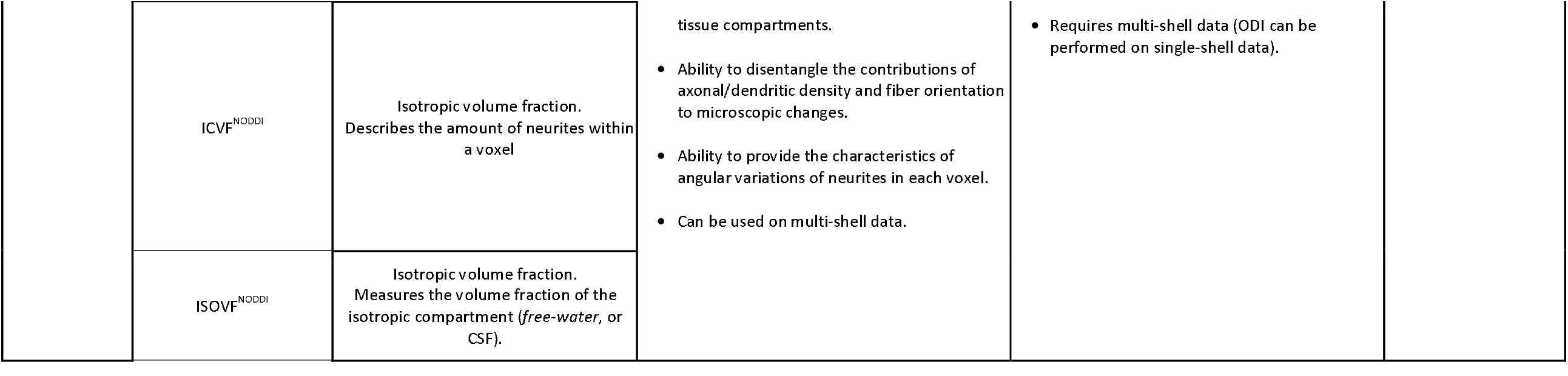
Diffusivity indices considered by this study. Diffusivity indices investigated in this study are described, alongside their advantages and limitations.

Main nuisance covariates included years of education, waist/hip ratio, population structure, measured using the first four genetic principal components, and the Townsend index. These covariates were also used in recent UK Biobank structural investigations (Lawrence et al. 2021; Salminen et al. 2022). In detail, educational attainment was assessed using a sociodemographic questionnaire in which participants reported their completed qualification level (UK education system). These results were generalized beyond the UK education system and converted to the corresponding number of years using ISCED harmonization guidelines (Goujon et al. 2016) and dichotomized into ‘high school’ versus ‘college’ (>=17 years) education. Waist-to-hip ratio is a common index of central fat distribution (Gadekar et al. 2020) and used as a proxy of atherosclerotic burden in overweight individuals and postmenopausal women (Lee et al. 2015; Scicali et al. 2018). Population structure was obtained from the UK Biobank’s genetic ancestry analyses (Bycroft et al. 2017). The first four genetic principal components were regressed out to account for differences in ethnicity within the sample. Participants’ socioeconomic status (SES) was measured using the Townsend Index (Townsend et al. 2012), a British measure of societal deprivation based on the postal code of each participant – this index has been often used as a proxy measure of SES (Smith et al. 2001). For any of the variables of interest, women who had missing data, responded ‘do not know’ or ‘preferred not to answer’ were excluded from each analysis.

Our primary hypotheses include a hormone by age interaction on advanced diffusivity metrics (ISOVF^NODDI^, OD^NODDI^, and ICVF^NODDI^). A previous investigation of age and sex effects on microstructure in the UK Biobank reported the highest effect sizes for advanced NODDI metrics in sex by age interactions on full white matter (Lawrence et al. 2021). False discovery rate (FDR) was used to account for multiple comparisons (corrected at *p*<.05) (Benjamini and Hochberg 1995). Secondary hypotheses include age by treatment type interaction on the advanced diffusivity NODDI metrics and a steeper aging trajectory for women taking unopposed estrogen treatment relative to combination estrogen users; additionally, a significant association between duration of hormone use, age at therapy onset and age at menopause for women taking hormones.

*Post-hoc* exploratory investigations considered the effects of HT compound composition (combined vs unopposed estrogen treatment) on WM aging trajectories; data was available for only a subset of the UK Biobank participants in the sample (unopposed estrogen users, n=300; combination users, n=98).

Supplemental analyses included Spearman’s correlations to investigate a potential association between significant dMRI measures and duration of HT and OC usage, age at which the participant began HT or OC therapy, and age at menopause in HT and OC users. For any significant association, these variables were additionally covaried for in our fractional polynomial analyses.

Furthermore, centile curves (Bethlehem et al. 2018; Nobis et al. 2019; Lv et al. 2020) for each diffusion metric with respect to age were calculated and plotted in R; kernel density plots, indicating the degree of data point overlap (and sampling density across the age range), are included in the plots.

### MRI processing

For detailed descriptions of the UK Biobank data acquisition, neuroimaging protocol, and validation, see work by Miller (Miller et al. 2016b) and Alfaro-Almagro and colleagues (Alfaro-Almagro et al. 2018). In brief, dMR images were acquired at b=1000 and 2000 s/mm^2^ along with 50 non-coplanar diffusion directions per shell with 5 b=0 s/mm^2^ (and 3 b=0 blip-reversed) using a standard (‘monopolar’) Stejskal-Tanner pulse sequence (voxel dimensions: (2 mm)^3^ isotropic, field of view: 104×104×72 mm; gradient timings: small delta=21.4 ms, big delta=45.5 ms) for a total imaging time of 7 minutes. All images underwent pre-processing, involving noise correction (Veraart et al. 2016), Gibbs-ringing correction (Kellner et al. 2016), estimation of echo-planar imaging distortions, motion and eddy current and susceptibility distortion corrections (Andersson and Sotiropoulos 2016), spatial smoothing *(fslmaths* in FSL) with Gaussian kernel (1 mm FWHM)^3^ and diffusion metric estimation. Diffusion tensor (DTI) and Neurite Orientation Dispersion and Density Imaging (NODDI) diffusion maps (Zhang et al. 2012) were computed by UK Biobank, while the tensor distribution function (TDF) (Leow et al. 2009; Nir, Jahanshad, Villalon-Reina, Isaev, Zavaliangos-Petropulu, Zhan, Leow, Jack Jr, et al. 2017) was computed locally using code available at https://git.ini.usc.edu/ibagari/TDF. Diffusivity indices considered by this study are detailed below and summarized in ***Table 3***. For more details on these methods, please refer to the *Supplemental Material*.

Diffusion tensor imaging (DTI) fitting was performed on the b=1000 s/mm^2^ shell (50 diffusion-encoding directions) using the DTI fitting (*DTIFIT*, FSL) tool to create maps of the following tensor-derived measures: fractional anisotropy (FA^DTI^), representing the degree of anisotropic diffusivity - often considered to be sensitive to the density of white matter fibers and the degree of directional coherence within a fiber bundle (Pierpaoli et al. 1996; Jones and Cercignani 2010). Other DTI indices included mean diffusivity (MD^DTI^), describing the molecular diffusion rate in a fiber bundle, radial diffusivity (RD^DTI^), reflecting diffusivity perpendicular to the axonal fibers (or the principal direction of diffusion), and axial diffusivity (AD^DTI^) describing the magnitude of diffusion parallel to fiber tracts.

Diffusion MRI data (b=1000 s/mm^2^, 50 diffusion-encoding directions) was also used as input data to estimate the tensor distribution function (TDF) at each voxel in the (Leow et al. 2009; Nir, Jahanshad, Villalon-Reina, Isaev, Zavaliangos-Petropulu, Zhan, Leow, Jack Jr, et al. 2017). This approach extends multi-tensor models of diffusion to describe intravoxel fibers mathematically as a probabilistic collection of tensors, or, alternatively, a continuous mixture of Gaussian densities. The fitted TDF function was employed to create diffusion maps of FA, based on the mixture of tensors (FA^TDF^). This approach overcomes some known limitations of the single diffusion tensor model in regions of fiber mixing or crossing and has been shown in other datasets to boost effect sizes for associations with external variables of interest, such as age, Alzheimer’s disease, or clinical measures of dementia severity (Nir, Jahanshad, Villalon-Reina, Isaev, Zavaliangos-Petropulu, Zhan, Leow, Jack Jr, et al. 2017; Villalon-Reina et al. 2018; Lawrence et al. 2021).

In addition to TDF and DTI models, the dMRI data (all shells) were input into Neurite Orientation Dispersion and Density Imaging (NODDI) (Zhang et al. 2012) biophysical models, using the AMICO (Accelerated Microstructure Imaging via Convex Optimization) tool (Daducci et al. 2015), to obtain the following voxel-wise microstructural parameters: ODI^NODDI^ (orientation dispersion index, a measure of within-voxel white matter tract disorganization), ICVF^NODDI^ (intracellular volume fraction, and index of white matter neurite density) and ISOVF^NODDI^ (isotropic or free water volume fraction). These metrics have been shown to be sensitive to aging effects in smaller samples than that analyzed here (Thomopoulos et al. 2020), including measures of brain amyloid burden measured with PET (Thomopoulos et al. 2021).

Voxel-wise statistical analysis of the images was performed using Tract-based Spatial Statistics (Smith et al. 2006) (TBSS; ***Figure 1***) using the <u>ENIGMA-DTI processing pipeline</u>, with the following steps: FA^DTI^ images were non linearly registered to a standard-space white matter skeleton, the ENIGMA FA^DTI^ white matter template, representing the center of all white matter FA voxels in MNI space using ANTs symmetric image normalization (SyN) method (Avants et al. 2009). The resulting transformation was applied to the maps of each metric (TDF and NODDI). White matter metrics were projected onto the ENIGMA template skeleton based on the FA^DTI^ metric using FSL’s TBSS. Average whole-brain measures of FA^DTI^, MD^DTI^, RD^DTI^, AD^DTI^, FA^TDF^, ODI^NODDI^, ISOVF^NODDI^, ICVF^NODDI^ were extracted for each participant. As part of the quality control processes, 2D images of the output from each processing step in the dMRI pipeline were visually reviewed. Careful visual checks were performed for individuals whose derived data were flagged as outliers. Prior to any group-level analysis, participants with low-quality data (such as gross visual artifacts) were eliminated. A total of N=114 subjects were excluded from our analyses.

**Figure 1.**
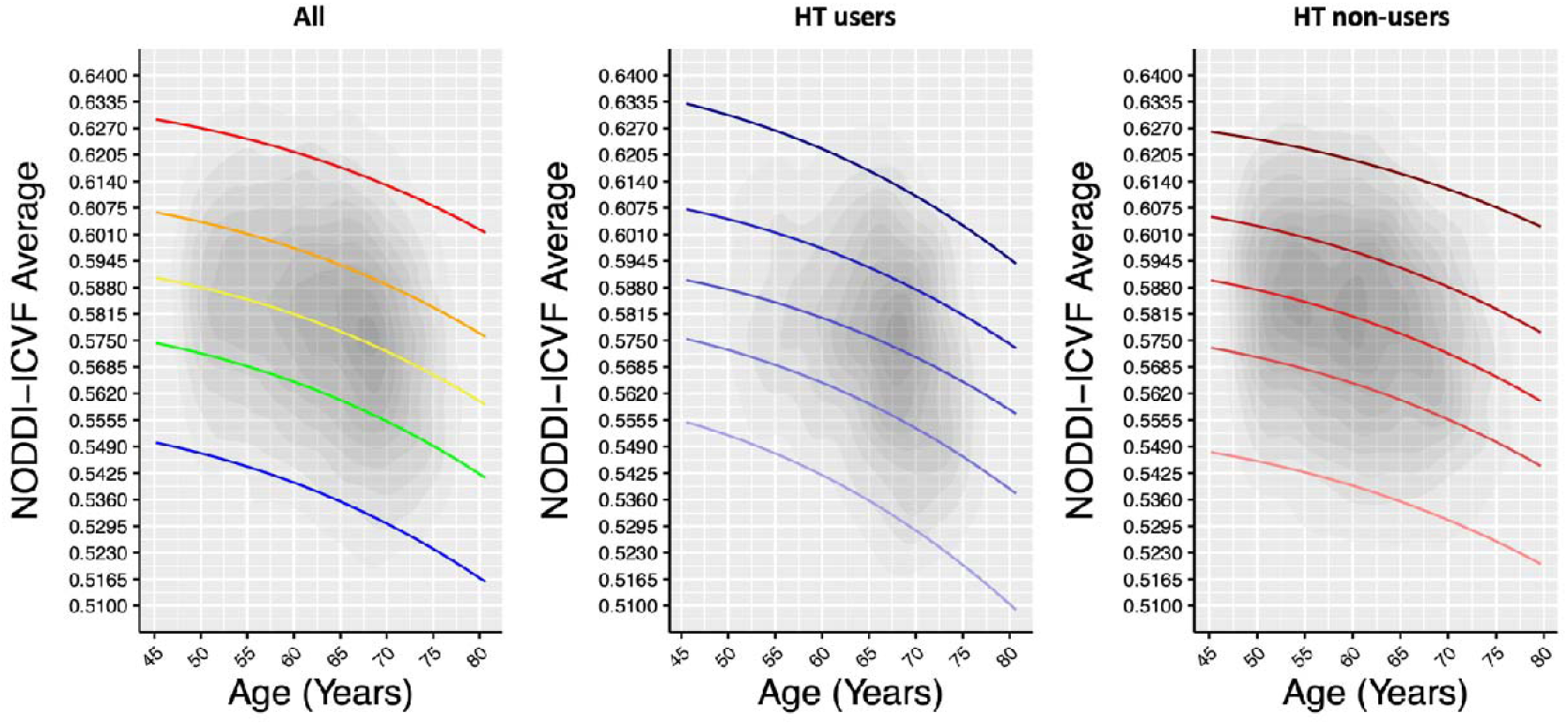
Normative centile reference curves for ICVF^NODDI^ in HT users and non-users. Whole brain white matter averages for ICVF^NODDI^ are shown for HT (*middle panel)* and non-users (*right panel*), and collectively (*left panel*). Solid colored lines, bottom-up, indicate the following centiles: 5th, 25th, 50th, 75th, 95th. Kernel densities indicating the degree of data point overlap (and sampling density across the age range), are shown in grey.

## RESULTS

### Age-related trajectories in HT

There was a significant interaction between age and HT status on diffusion metrics ICVF^NODDI^; (decreased intra-cellular volume fraction with age; p=.0009; *age*^*(p0*.*5)*^) when including the main nuisance covariates (**Figure 1**). We did not detect significant interactions between age and HT status on diffusion metrics DTI or TDF. Age effects on ICVF^NODDI^ were confirmed when adjusting for history of OC (decreased with age; p=.001; *age*^*(p0*.*5)*^). Supplemental analyses also adjusting for surgical history (hysterectomy and oophorectomy), cancer history, and number of childbirths as nuisance covariates did not confirm the significant age effect seen on diffusivity metric ICVF^NODDI^ in HT users and non-users. Within the group of HT users (N=3,033), no association was found between the significant dMRI measures and duration of HT use, age at HT initiation and age at menopause (all p>.05).

Follow-up *post-hoc* exploratory analyses in the HT users group – examining unopposed (N=300, age (mean±SD)=61.1±7.4) and combined (N=90; age (mean±SD)=58.4±6.2) estrogen treatments – revealed a significant age interaction on diffusion metric ISOVF^NODDI^ (increased free water with age in both groups; p=.033; *age*^*(p1)*^; ***Figure 2***). No significant interaction was found on diffusion metrics DTI or TDF. Adjusting for history of OC confirmed findings on ISOVF^NODDI^ (increased with age; p=.034; *age*^*(p1)*^). Additionally adjusting for duration of HT, age at HT onset and age at menopause, did not alter age effects seen on ISOVF^NODDI^ in HT users using unopposed and combination estrogen treatments (p=.01; *age*^*(p1)*^, p=.01; *age*^*(p1)*^, and p=.02; *age*^*(p1)*^, respectively). Specifically, in combination HT (estrogen + progestin) users, later age at HT onset (*left panel* **Figure 1 *Supp Material***), prolonged duration of HT therapy (*middle panel*), and later age at menopause (*right panel*), were associated with higher ISOVF^NODDI^, relative to estrogen only HT users. In the estrogen only HT group, no significant association was found between clinical variables duration of use, age at initiation and age at menopause and ISOVF^NODDI^ (r=0 to 0.1, p=.9 to 1.0). In the combination HT group, a weak positive correlation was found between duration of HT and ISOVF^NODDI^ (r=.2, p=.04), while no significant association was found between ISOVF^NODDI^ and age at HT initiation (r=.15, p=.16), or age at menopause (r=.18, p=.13) in this group (***Figure 1 Supp Material***).

**Figure 2.**
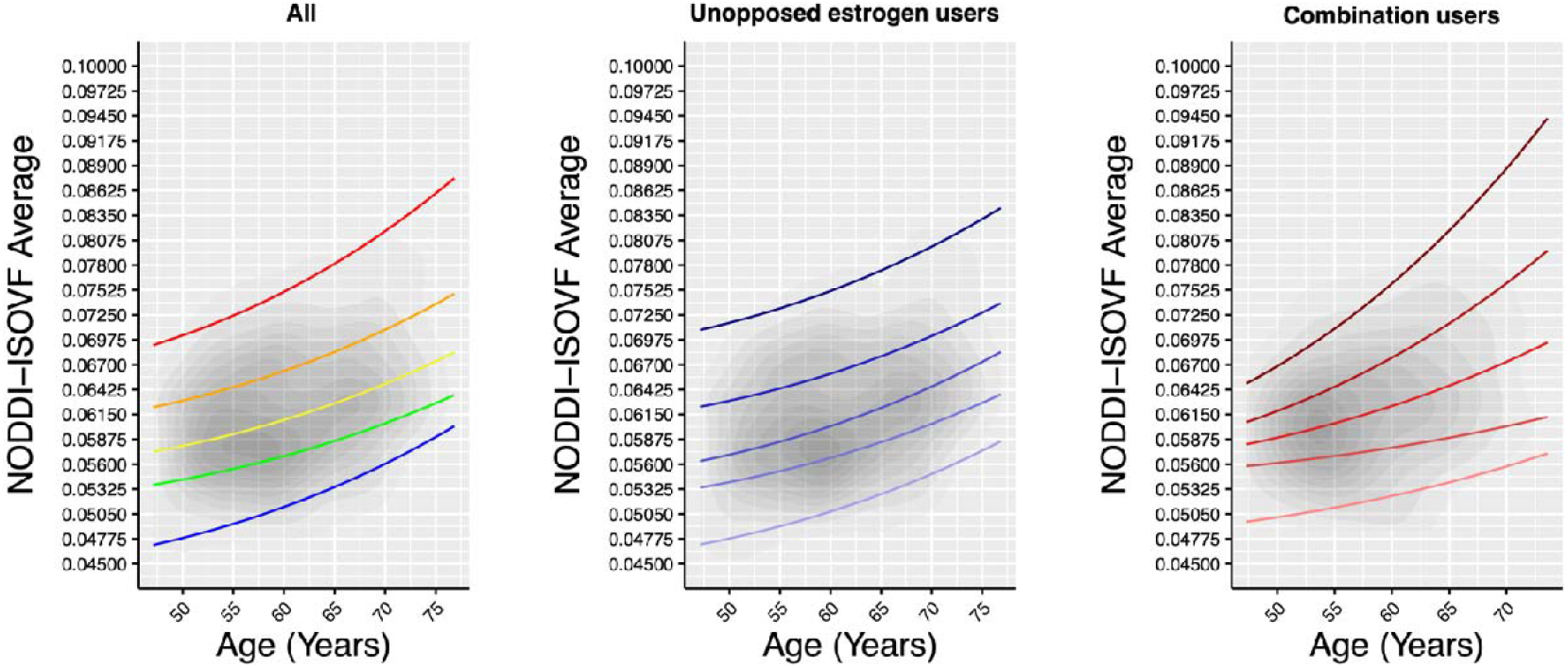
Normative centile reference curves for ISOVF^NODDI^ in unopposed and combination estrogen users. Whole brain white matter averages for ISOVF^NODDI^ shown for estrogen users and combination users, collectively (*left panel*) and separately for unopposed estrogen (*middle panel*) and combination (*right panel*) users. Unopposed estrogen users (N=300) displayed a steeper ISOVF^NODDI^ aging trajectory compared to combination users (N=98). Solid colored lines, bottom-up, indicate the following centiles: 5th, 25th, 50th, 75th, 95th. Kernel densities indicating the degree of data point overlap (and sampling density across the age range), are shown in grey.

Representative centile curves for each significant diffusion metric are presented in ***Figures 1 and 2*** with colored lines indicating 5th (black), 25th (blue), 50th (red), 75th (green), and 95th (purple) centiles, and kernel density maps indicating the degree of data point overlap. Curves show age-dependent trajectories for ICVF^NODDI^ (***Figure 1***), and ISOVF^NODDI^ (***Figure 2***). In summary, all the HT status effects on the diffusion metrics were in the direction of ‘more aging’ – decreased ICVF^NODDI^, and increased ISOVF^NODDI^.

### Age-related trajectories in OC

There was no significant interaction between age and OC status on any of the diffusion metrics assessed when including the main nuisance covariates (p>.05). Supplemental analyses including surgical history (hysterectomy and oophorectomy), cancer history, and number of childbirths as nuisance covariates confirmed no significant interaction between diffusion metrics and age in OC users and non-users (p>.05).

## DISCUSSION

We examined the association between exogenous sex hormone exposure and the brain’s WM aging trajectories in post-menopausal middle aged and older women with and without hormone therapy (HT), as well as oral contraceptives (OC). To thoroughly examine microstructure, we used 3 increasingly sophisticated diffusion models, from single-shell tensor-based (DTI and TDF) to multi-shell (NODDI) methods. We also present normative reference curves for various aspects of white matter microarchitecture in women, which may one day allow us to identify individuals with the most severe neural pathology.

As a whole, age and sex hormone exposure usage were both statistically associated with WM metrics, with advanced dMRI model NODDI capturing these differences most sensitively. We found that HT usage was associated with less favorable aging patterns, observed as more pronounced white matter changes on advanced white matter metrics, relative to those who did not use HT (**Figure 1**). The observed less favorable aging trajectories in this sample were independent of previous OC usage; however, history of pregnancies, surgeries, or cancer were found to influence these trajectories. Furthermore, accounting for differences in hormone therapy formulation, we showed that not all HT formulations may exert a detectable neuroprotective effect, at least on the white matter measures we examined here. Hormone replacement therapy containing conjugated estrogens may be associated with marginally more beneficial aging patterns (less pronounced white matter changes) than formulations containing estrogen alone (**Figure 2**). These aging effects on white matter metrics were independent of the duration of hormone therapy, the age at which hormone therapy was initiated and the age at which women entered menopause (**Figure 1 Supp**). Moreover, OC was not found to alter the derived white matter measures considered in this study, nor did clinical factors modulate white matter aging trajectories in OC users and non-users. To the best of our knowledge, the current work is the first study of the associations between exogenous hormone exposure effects on aging in white matter in a population-based cohort.

HT usage was associated with a steeper WM aging trajectory, and this was evidenced by lower neurite density (ICVF^NODDI^) in the HT user group relative to non-users (**Figure 1**). Furthermore, unopposed estrogen usage was associated with a steeper WM aging trajectory, and this was evidenced by increased isotropic volume fraction (ISOVF^NODDI^) relative to combination estrogen usage. NODDI is a multi-shell model of tissue microstructure developed to better describe the underlying WM tissue structure than traditional diffusion-tensor models like DTI or its mathematical extension TDF. While tensor-based models assume that each voxel comprises of a single tissue compartment, NODDI directly models multiple aspects of the cellular environment (Zhang et al., 2012). Notably, the NODDI model can remove the effect of CSF contamination (e.g., following cortical atrophy due to aging processes), increasing the specificity to the cytoarchitecture subjected to investigation. The diffusion metric ISOVF^NODDI^ represents the fraction of water molecules that are not restricted or directed and is also described as ‘free water’. Higher free water may also be an index of neuroinflammation as neuroinflammatory processes can increase the proportion of water molecules diffusing freely in the extracellular space. Thus, the observed increase in circulating sex-hormone levels following HT use in postmenopausal women, and specifically in women taking HT formulations containing estrogen alone, may suggest increased susceptibility to mechanisms of neuroinflammation or neuronal loss in this group. These findings are in line with our hypothesis that sex hormone therapy would *negatively* impact women’s WM aging. This corroborates prior reports of negative effects of exogenous sex hormones on brain structural measures and expands them to include WM microstructural deficits. Prior aging and AD research has implicated the progression of tau protein to neurofibrillary tangles and the resulting destabilization of axonal microtubules (Spillantini and Goedert 2013), as well as cortical grey matter volumetric reductions and neuronal dispersion (Colgan et al. 2016). It is possible that in postmenopausal women, with age, as the cortical structure atrophies, the space becomes occupied by CSF (or free water), which is reflected by higher isotropic volume fraction values. While further clinical measures and investigations at the microstructural level are necessary, our findings may suggest increased susceptibility to developing tau-dependent pathologies such as AD later in life in middle- and older-aged women with a history of HT use, specifically in estrogen-only HT users. Conversely, our findings are consistent with a neuroprotective effect of sex hormones during normal development but a subsequent deterioration in WM metrics in later life in women using combination HT.

We failed to detect changes in WM aging trajectories in middle- and older-aged women following OC treatment. Prior investigations in the UK Biobank reported reductions in cortical gray matter volume in OC users relative to never-users, worsening by duration of OC treatment (de Lange et al. 2019). To the best of our knowledge, this is the largest investigation of OC effects on white matter aging trajectories in middle-aged to older women. While it is possible that chronic ovarian hormone suppression (induced by OC treatment) may progressively affect WM microstructure (as well as cortical volume) later in life, longitudinal studies are needed to draw further conclusions. We cannot exclude that the lack of detectable effects in OC users on the brain’s WM in this sample may depend on the type of OC formulation, which was not possible to assess in the current study with the currently available data in the UK Biobank. Birth control formulations vary from ethinylestradiol-only formulations to those with added concentrations of progestin, and we may expect that differences in formulation may exert a differential effect on diffusion metrics. With this in mind, future studies should disentangle effects of birth control formulation on aging trajectories in women. There are also changes occurring in the societal use and duration and timing of OC use. The number of women availing of OC treatment is increasing, while the age of first contraceptive use is constantly decreasing to sensitive neuroplastic periods during puberty. Given these factors, future research efforts are warranted to investigate OC effects on the brain longitudinally across the lifespan.

The female immune system changes significantly during pregnancy and menopause (Mor et al. 2011; Mishra and Brinton 2018), and evidence suggests that immune regulation during these major transitional phases may influence women’s brain aging trajectories later in life (Ding et al. 2013; Mosconi et al. 2017; Fox et al. 2018). Hormonal fluctuations and their interactions with immune processes play a role in maternal brain adaptations, but their long-term effects on brain aging in women are unknown (Barth and de Lange 2020). Pregnancies are associated with alterations in microglia density, number, and activity which could modulate neuroplastic compensation in response to perimenopausal inflammation processes (Barth and de Lange 2020). The E2 receptor is a key neuroplasticity regulator in the female brain (Barha and Galea 2010), and it has been suggested that neuroplasticity may be important in understanding the boundaries between normal aging and the early stages of Alzheimer’s disease (Fjell et al. 2014). In our analyses, the number of childbirths may modulate the age effects on neurite density (a steeper aging WM trajectory) in HT users relative to non-users. Pregnancy-related immune adaptations, in conjunction with endocrinological modulations, may influence maternal brain plasticity during pregnancy and *post-partum*, potentially influencing the course of neurobiological aging later in life. In this study women had on average ∼2 children, thus we cannot exclude that a higher number of childbirths, alongside other clinically relevant factors (such as number of abortions and miscarriages), may differentially influence aging trajectories on WM metrics. Prior investigations within the UK Biobank have shown that subjects with two or three offspring had significantly reduced brain age compared to those without offspring (Ning et al., 2020). Although beyond the scope of this analysis, irrespective of HT, we found that parity affected the dispersion of intra-voxel freely diffusing water molecules (ISOVF^NODDI^) overall in the brain, which may corroborate our study findings and findings by Ning et al., (2020) on apparently reduced brain aging and altered cognitive function in subjects with two or three offspring (Ning et al. 2020).

The ‘healthy cell bias of estrogen action hypothesis’ (Brinton 2008) suggests that neuronal viability and general health - before starting hormone therapy - might be of importance for exogenous estrogens to exert therapeutic effects. This hypothesis may be relevant for women undergoing surgeries that can cause a drop in circulating estrogen levels such as hysterectomy and/or oophorectomy and lead to early menopause. A history of hysterectomy and/or oophorectomy was accounted for in the present study and was found to influence age effects on neurite density seen in HT users relative to non-users. Our results support the ‘critical period hypothesis’ for estrogens, as an early initiation of (combination) HT was associated with less evident white matter aging (particularly before menopause, as findings did not change when covarying for age at menopause). Of note, HT can be also used to slow or stop the growth of cancers that use hormones to grow, such as some breast cancers. In this study, age effects on neurite density seen in HT users relative to non-users were influenced by cancer history in women (also including breast, womb, and ovarian cancers).

The second theory regarding hormone therapy effects is known as the ‘critical period hypothesis’ (Maki 2013), whereby HT may be neuroprotective if initiated soon after cessation of ovarian function (<5 years from menopause; (MacLennan et al. 2006; Erickson et al. 2010; Espeland et al. 2015), but carry detrimental effects if initiated much later (>10 years from menopause, (Hodis et al. 2016). Substantial evidence suggests that HT formulation, administration, dosage, compound composition, and mode of administration are clinically relevant to the course of aging in women (Gleason et al. 2015; Savolainen-Peltonen et al. 2019). Within the groups of HT and OC users, we failed to detect any meaningful association between the significant dMRI measures and duration of HT use, age at HT initiation and age at menopause (all p>0.05). We observed a weak positive association between duration of HT and ISOVF^NODDI^, whereby later commencement of therapy was associated with higher volume fraction of extracellular isotropic free water in the combination estrogen group (trends depicted in **Figure 1 Supp**). Even so, controlling for duration of treatment in our analyses did not alter age effects on ISOVF^NODDI^ in this treatment group. We cannot exclude that later commencement of HT may be associated with greater reductions in age-dependent metrics of WM microstructure beyond those assessed by the present study, and that other clinical factors (e.g., examining genotype interactions) not considered by the present study may further explain this association. Observing the effects of HT on brain aging in women within such a short interval (within 5 years from menopause) could have implications for understanding hormone-related dementia pathogenesis in women. Future studies should account for such differences to disentangle effects of HT treatment regime on WM microstructural decline in women and identify other vulnerability factors and critical age windows for administering HT in women.

Estrogens interact with multiple neurotransmitter systems in our brain to modulate mood and cognition (Barth et al. 2015); in particular, the interaction with dopamine might be of interest considering its role in pathophysiological processes of neurological, as well as psychiatric disorders, and its modulation of executive functions such as working memory and reward processing which are often impaired in subjects with dementia or AD (Reeves et al. 2009). Estrogen receptors (ERs) - ERα and Erβ – are widely distributed in the brain and found in subcortical limbic gray matter and basal ganglia areas, as well as prefrontal and temporal cortices (Barth et al. 2015). Studies focusing on the role of estrogen receptors and endogenous estrogen signaling have reported neuroprotective effects on the brain, via their effect promoting synthesis of neurotrophins and protecting the brain from inflammation and stress (Brann et al. 2007). The expression of the ERs can be overlapping or distinct, dependent upon brain region, sex, age, and exposure to hormones. During the time of menopause, there may be changes in receptor expression profiles, post-translational modifications, and protein-to-protein interactions that could lead to a very different environment for estrogen to exert its effects (Mott and Pak 2013). These effects have been considered in clinical trials of those suffering from psychiatric disorders where estrogen-signaling pathways have been shown to be compromised (Hwang et al. 2021). Changes in circulating estrogen levels may modulate the microstructure of WM and contribute to pathophysiological processes of neurological and psychiatric disorders in women with a history of HT.

Moreover, studies in experimental animal models have established a role for sex hormones including estrogen in WM abnormalities and have provided a convincing rationale for HT in prevention and treatment of dementia (Birge 1996; Cerghet et al. 2009). Interventional trials of estrogen and its analogs (often in conjunction with other adjuvants) are ongoing to investigate the long-term safety and efficacy of these compounds to restore structural integrity and related cognitive function of the brain in participants with dementia and early-stage AD. A comprehensive and integrated understanding of estrogens and estrogen signaling across multiple levels of the brain system architecture, from cellular to molecular to systemic, is key to elucidate the mechanisms involved in its effects in diseased brains. Future studies are warranted to examine longitudinal white matter changes in women following sex-hormone therapy – both HT and OC.

There are some methodological considerations in this study. Besides using multiple statistical approaches, a strength of the current work was the inclusion of multiple diffusion metrics, from tensor-based to “beyond-tensor” models to provide a richer understanding of the underlying WM structure than traditional diffusion metrics alone. Besides the tensor-based metric, FA, we considered the estimated number of crossing fibers, fiber complexity, neurite dispersion and estimated numbers of fiber compartments. Diffusion metric FA can lack specificity as it does not directly delineate changes in tissue microstructure – for example a reduction in FA can be associated with different types of microstructural changes, such as demyelination, or a reduction in axonal density. A decline in FA with age could result from many cellular factors, including decreases in the density or an increase in the dispersion orientation of neurites (Pines et al. 2020). Multi-compartment models such as NODDI attempt to overcome this by modeling diffusion data using a set of indices that are more directly related to WM microstructure.

Moreover, TBSS is a widely used method for whole-brain voxel-based analysis but has some limitations. TBSS reduces individual white matter tracts to a skeleton, delineating the center of the tracts and projecting onto it only the highest value of FA along the projection. This may result in loss of microstructural information and potential artifacts (Bach et al. 2014), some due to misregistration. In this study, we carried out careful qualitative and quantitative assessments of registration for each subject’s diffusion metric. Additionally, while we measured diffusivity indices on the FA skeleton, some of these indices may not have a local maximum at the exact same location as FA. Future studies should investigate if findings hold without white matter skeletonization, i.e., without warping images into a predefined template, but by averaging values within regions of interest (ROIs) in native space (Nir et al. 2021). TBSS has limited anatomical specificity (adjacent tracts may not be distinguished) and thus may be susceptible to false positives (Bach et al. 2014). Also, while these limitations may arise for tract-specific diffusion metrics, we chose to focus our analyses on whole-brain diffusion averages.

Unlike a randomized clinical trial, where treatments are randomized to individuals, and the effects are studied longitudinally, the current design is a naturalistic study of people using medications. As such, we report associations that may still be important in understanding brain aging trends in a broad and inclusive population and suggest mechanisms that could be tested in a causally informed design (such as an RCT). To further understand confounders in the population, future studies could seek demographic, educational, or cultural factors that might affect HT/OC use or access, and test if such factors might mediate any detected effects.

We also note that the meaning of the term “trajectories” may differ when inferred from cross-sectional rather than longitudinal data, as individuals at different parts of the age range may differ from each other in their childhood developmental experiences, environments, and health risks throughout life. As the participants range in age by almost 40 years, any interactions with age could also be due to societal changes in the incidence of use, duration of use, and breadth and diversity of access to HTs and OCs. Also, attrition of individuals late in life (due to mortality, or ill health) means that older individuals who participate in the study may show less steep aging trajectories than a typical individual in the population (due to selection bias, or survivor bias).

This study shows the value of testing alternative models for lifespan trajectories beyond popular linear and quadratic models, especially when dealing with large samples. The fractional polynomial approach may offer a more flexible alternative to linear or quadratic models and may be useful for estimating age effects on health outcomes. Large scale datasets such as the UK Biobank offer the opportunity to test alternative models for lifespan trajectories beyond traditional statistical models. Studies with a large number of data points may benefit from evaluating findings across alternative explanatory models to understand the robustness of findings.

The cross-sectional nature of this study does not enable causal inference, so future studies should use longitudinal designs – ideally randomized – which are vital to fully understand how estrogen exposure influences brain aging across the lifespan. The various effects of exogenous sex hormones on brain function deserve greater critical observation to unravel possible pathogenetic mechanisms, as well as to identify individuals at high risk of hormone therapy-related adverse consequences such as dementia or AD. Future studies may benefit from including further clinical and plasma measures and examining genotype interactions (e.g., with apolipoprotein E genotype) to better understand how estrogen exposure influences brain aging and genetic predisposition for neurological disorders in women. Such studies have long-term public health implications and may eventually improve early detection, clinical intervention, and quality of life for individuals at risk for age-related neurodegenerative disorders.

## Acknowledgments

This research was conducted using the UK Biobank dataset via an approved application, #11559. Our study was supported by NIH grants R01 AG060610, R01AG058854, U54 EB020403, U01AG068057, R01 AG059874, and P41 EB015922.

## REFERENCES

Alfaro-Almagro F, Jenkinson M, Bangerter NK, Andersson JLR, Griffanti L, Douaud G, Sotiropoulos SN, Jbabdi S, Hernandez-Fernandez M, Vallee E. 2018. Image processing and Quality Control for the first 10,000 brain imaging datasets from UK Biobank. Neuroimage. 166:400–424.

Andersson JLR, Sotiropoulos SN. 2016. An integrated approach to correction for off-resonance effects and subject movement in diffusion MR imaging. Neuroimage. 125:1063–1078.

Avants BB, Tustison N, Song G. 2009. Advanced normalization tools (ANTS). Insight j. 2:1–35.

Bach M, Laun FB, Leemans A, Tax CMW, Biessels GJ, Stieltjes B, Maier-Hein KH. 2014. Methodological considerations on tract-based spatial statistics (TBSS). Neuroimage. 100:358–369.

Barha CK, Galea LAM. 2010. Influence of different estrogens on neuroplasticity and cognition in the hippocampus. Biochimica et Biophysica Acta (BBA)-General Subjects. 1800:1056–1067.

Barth C, de Lange AMG. 2020. Towards an understanding of women’s brain aging: the immunology of pregnancy and menopause. Front Neuroendocrinol. 58:100850.

Barth C, Steele CJ, Mueller K, Rekkas VP, Arélin K, Pampel A, Burmann I, Kratzsch J, Villringer A, Sacher J. 2016. In-vivo Dynamics of the Human Hippocampus across the Menstrual Cycle. Sci Rep. 6:1–9.

Barth C, Villringer A, Sacher J. 2015. Sex hormones affect neurotransmitters and shape the adult female brain during hormonal transition periods. Front Neurosci. 9:37.

Basser PJ, Mattiello J, LeBihan D. 1994. MR diffusion tensor spectroscopy and imaging. Biophys J. 66:259–267.

Benjamini Y, Hochberg Y. 1995. Controlling the false discovery rate: a practical and powerful approach to multiple testing. Journal of the Royal statistical society: series B (Methodological). 57:289–300.

Bennett IJ, Madden DJ. 2014. Disconnected aging: cerebral white matter integrity and age-related differences in cognition. Neuroscience. 276:187–205.

Bethlehem RAI, Seidlitz J, Romero-Garcia R, Dumas G, Lombardo M V. 2018. Normative age modelling of cortical thickness in autistic males. bioRxiv.

Birge SJ. 1996. Is there a role for estrogen replacement therapy in the prevention and treatment of dementia? J Am Geriatr Soc. 44:865–870.

Boyle CP, Raji CA, Erickson KI, Lopez OL, Becker JT, Gach HM, Kuller LH, Longstreth W, Carmichael OT, Riedel BC, Thompson PM. 2020. Estrogen, brain structure, and cognition in postmenopausal women. Hum Brain Mapp. 1–12.

Brann DW, Dhandapani K, Wakade C, Mahesh VB, Khan MM. 2007. Neurotrophic and neuroprotective actions of estrogen: basic mechanisms and clinical implications. Steroids. 72:381–405.

Brinton RD. 2008. The healthy cell bias of estrogen action: mitochondrial bioenergetics and neurological implications. Trends Neurosci. 31:529–537.

Bycroft C, Freeman C, Petkova D, Band G, Elliott LT, Sharp K, Motyer A, Vukcevic D, Delaneau O, O’Connell J, Cortes A, Welsh S, McVean G, Leslie S, Donnelly P, Marchini J. 2017. Genome-wide genetic data on ∼500,000 UK Biobank participants. bioRxiv.

Cerghet M, Skoff RP, Swamydas M, Bessert D. 2009. Sexual dimorphism in the white matter of rodents. J Neurol Sci. 286:76–80.

Christin-Maitre S. 2013. History of oral contraceptive drugs and their use worldwide. Best Pract Res Clin Endocrinol Metab. 27:3–12.

Colgan N, Siow B, O’Callaghan JM, Harrison IF, Wells JA, Holmes HE, Ismail O, Richardson S, Alexander DC, Collins EC, Fisher EM, Johnson R, Schwarz AJ, Ahmed Z, O’Neill MJ, Murray TK, Zhang H, Lythgoe MF. 2016. Application of neurite orientation dispersion and density imaging (NODDI) to a tau pathology model of Alzheimer’s disease. Neuroimage. 125:739–744.

Costantino M, Pigeau G, Parent O, Ziolkowski J, Devenyi GA, Gervais NJ, Chakravarty MM. 2022. Menopause, Brain Anatomy, Cognition and Alzheimer’s Disease.

Covan EK. 2005. Meaning of aging in women’s lives. J Women Aging. 17:3–22.

Cox SR, Ritchie SJ, Tucker-Drob EM, Liewald DC, Hagenaars SP, Davies G, Wardlaw JM, Gale CR, Bastin ME, Deary IJ. 2016. Ageing and brain white matter structure in 3,513 UK Biobank participants. Nat Commun. 7:1–13.

Daducci A, Canales-Rodríguez EJ, Zhang H, Dyrby TB, Alexander DC, Thiran J-P. 2015. Accelerated microstructure imaging via convex optimization (AMICO) from diffusion MRI data. Neuroimage. 105:32–44.

Daniels K, Abma JC. 2019. Current contraceptive status among women aged 15–49: United States, 2015–2017. NCHS data brief no. 327. 2018.

de Lange AG, Barth C, Kaufmann T, Maximov II, van der Meer D, Agartz I, Westlye LT. 2020. Women’s brain aging: Effects of sex-hormone exposure, pregnancies, and genetic risk for Alzheimer’s disease. Hum Brain Mapp. 41:5141–5150.

de Lange A-M, Barth C, Kaufmann T, Maximov II, van der Meer D, Agartz I, Westlye LT. 2019. Cumulative estrogen exposure, APOE genotype, and women’s brain aging-a population-based neuroimaging study. bioRxiv. 826123.

den Heijer T, Geerlings MI, Hofman A, de Jong FH, Launer LJ, Pols HAP, Breteler MMB. 2003. Higher estrogen levels are not associated with larger hippocampi and better memory performance. Arch Neurol. 60:213–220.

Dima D, Modabbernia A, Papachristou E, Doucet GE, Agartz I, Aghajani M, Akudjedu TN, Albajes-Eizagirre A, Alnæs D, Alpert KI. 2022. Subcortical volumes across the lifespan: Data from 18,605 healthy individuals aged 3–90 years. Hum Brain Mapp. 43:452–469.

Ding M, Leach M, Bradley H. 2013. The effectiveness and safety of ginger for pregnancy-induced nausea and vomiting: a systematic review. Women and Birth. 26:e26–e30.

Erickson KI, Voss MW, Prakash RS, Chaddock L, Kramer AF. 2010. A cross-sectional study of hormone treatment and hippocampal volume in postmenopausal women: evidence for a limited window of opportunity. Neuropsychology. 24:68.

Espeland MA, Brinton RD, Hugenschmidt C, Manson JE, Craft S, Yaffe K, Weitlauf J, Vaughan L, Johnson KC, Padula CB. 2015. Impact of type 2 diabetes and postmenopausal hormone therapy on incidence of cognitive impairment in older women. Diabetes Care. 38:2316–2324.

Fjell AM, McEvoy L, Holland D, Dale AM, Walhovd KB, Initiative ADN. 2014. What is normal in normal aging? Effects of aging, amyloid and Alzheimer’s disease on the cerebral cortex and the hippocampus. Prog Neurobiol. 117:20–40.

Fox M, Berzuini C, Knapp LA, Glynn LM. 2018. Women’s pregnancy life history and Alzheimer’s risk: can immunoregulation explain the link? Am J Alzheimers Dis Other Demen. 33:516–526.

Frangou S, Modabbernia A, Williams SCR, Papachristou E, Doucet GE, Agartz I, Aghajani M, Akudjedu TN, Albajes-Eizagirre A, Alnæs D. 2022. Cortical thickness across the lifespan: Data from 17,075 healthy individuals aged 3–90 years. Hum Brain Mapp. 43:431–451.

Gadekar T, Dudeja P, Basu I, Vashisht S, Mukherji S. 2020. Correlation of visceral body fat with waist–hip ratio, waist circumference and body mass index in healthy adults: A cross sectional study. Med J Armed Forces India. 76:41–46.

Galea LAM, Leuner B, Slattery DA. 2014. Hippocampal plasticity during the peripartum period: influence of sex steroids, stress and ageing. J Neuroendocrinol. 26:641–648.

Gleason CE, Dowling NM, Wharton W, Manson JE, Miller VM, Atwood CS, Brinton EA, Cedars MI, Lobo RA, Merriam GR. 2015. Effects of hormone therapy on cognition and mood in recently postmenopausal women: findings from the randomized, controlled KEEPS–cognitive and affective study. PLoS Med. 12:e1001833.

Gogos A. 2013. Natural and synthetic sex hormones: effects on higher-order cognitive function and prepulse inhibition. Biol Psychol. 93:17–23.

Goujon A, Samir KC, Speringer M, Barakat B, Potancoková M, Eder J, Striessnig E, Bauer R, Lutz W. 2016. A harmonized dataset on global educational attainment between 1970 and 2060–An analytical window into recent trends and future prospects in human capital development. J Demogr Economics. 82:315–363.

Ha DM, Xu J, Janowsky JS. 2007. Preliminary evidence that long-term estrogen use reduces white matter loss in aging. Neurobiol Aging. 28:1936–1940.

Hodis HN, Mack WJ, Henderson VW, Shoupe D, Budoff MJ, Hwang-Levine J, Li Y, Feng M, Dustin L, Kono N. 2016. Vascular effects of early versus late postmenopausal treatment with estradiol. New England Journal of Medicine. 374:1221–1231.

Hoekzema E, Barba-Müller E, Pozzobon C, Picado M, Lucco F, García-García D, Soliva JC, Tobeña A, Desco M, Crone EA. 2017. Pregnancy leads to long-lasting changes in human brain structure. Nat Neurosci. 20:287–296.

Hwang WJ, Lee TY, Kim NS, Kwon JS. 2021. The Role of Estrogen Receptors and Their Signaling across Psychiatric Disorders.

Isaev DY, Nir TM, Jahanshad N, Villalon-Reina JE, Zhan L, Leow AD, Thompson PM. 2017. Improved clinical diffusion MRI reliability using a tensor distribution function compared to a single tensor. In: 12th International Symposium on Medical Information Processing and Analysis. SPIE. p. 464–471.

Jones DK. 2008. Studying connections in the living human brain with diffusion MRI. Cortex. 44:936–952.

Jones DK, Cercignani M. 2010. Twenty-five pitfalls in the analysis of diffusion MRI data. NMR Biomed. 23:803–820.

Kantarci K, Tosakulwong N, Lesnick TG, Zuk SM, Gunter JL, Gleason CE, Wharton W, Dowling NM, Vemuri P, Senjem ML. 2016. Effects of hormone therapy on brain structure: a randomized controlled trial. Neurology. 87:887–896.

Kellner E, Dhital B, Kiselev VG, Reisert M. 2016. Gibbs-ringing artifact removal based on local subvoxel-shifts. Magn Reson Med. 76:1574–1581.

Lawrence KE, Nabulsi L, Santhalingam V, Abaryan Z, Villalon-Reina JE, Nir TM, Ba Gari I, Zhu AH, Haddad E, Muir AM, Laltoo E, Jahanshad N, Thompson PM. 2021. Age and sex effects on advanced white matter microstructure measures in 15,628 older adults: A UK biobank study. Brain Imaging Behav.

Lee HJ, Hwang SY, Hong HC, Ryu JY, Seo JA, Kim SG, Kim NH, Choi DS, Baik SH, Choi KM. 2015. Waist-to-hip ratio is better at predicting subclinical atherosclerosis than body mass index and waist circumference in postmenopausal women. Maturitas. 80:323–328.

Leow AD, Zhu S, Zhan L, McMahon K, de Zubicaray GI, Meredith M, Wright MJ, Toga AW, Thompson PM. 2009. The tensor distribution function. Magn Reson Med. 61:205–214.

Lisofsky N, Mårtensson J, Eckert A, Lindenberger U, Gallinat J, Kühn S. 2015. Hippocampal volume and functional connectivity changes during the female menstrual cycle. Neuroimage. 118:154–162.

Lv J, Biase M Di, Cash RFH, Cocchi L, Cropley V, Klauser P, Tian Y, Bayer J, Schmaal L, Cetin-Karayumak S, Rathi Y, Pasternak O, Bousman C, Pantelis C, Calamante F, Zalesky A. 2020. Individual deviations from normative models of brain structure in a large cross-sectional schizophrenia cohort. bioRxiv. 2020.01.17.911032.

MacLennan AH, Henderson VW, Paine BJ, Mathias J, Ramsay EN, Ryan P, Stocks NP, Taylor AW. 2006. Hormone therapy, timing of initiation, and cognition in women aged older than 60 years: the REMEMBER pilot study. Menopause. 13:28–36.

Maki PM. 2013. The critical window hypothesis of hormone therapy and cognition: a scientific update on clinical studies. Menopause. 20:695.

Miller KL, Alfaro-Almagro F, Bangerter NK, Thomas DL, Yacoub E, Xu J, Bartsch AJ, Jbabdi S, Sotiropoulos SN, Andersson JLR, Griffanti L, Douaud G, Okell TW, Weale P, Dragonu I, Garratt S, Hudson S, Collins R, Jenkinson M, Matthews PM, Smith SM. 2016a. Multimodal population brain imaging in the UK Biobank prospective epidemiological study. Nat Neurosci. 19:1523–1536.

Miller KL, Alfaro-Almagro F, Bangerter NK, Thomas DL, Yacoub E, Xu J, Bartsch AJ, Jbabdi S, Sotiropoulos SN, Andersson JLR, Griffanti L, Douaud G, Okell TW, Weale P, Dragonu I, Garratt S, Hudson S, Collins R, Jenkinson M, Matthews PM, Smith SM. 2016b. Multimodal population brain imaging in the UK Biobank prospective epidemiological study. Nat Neurosci. 19:1523–1536.

Mishra A, Brinton RD. 2018. Inflammation: bridging age, menopause and APOEε4 genotype to Alzheimer’s disease. Front Aging Neurosci. 10:312.

Mor G, Cardenas I, Abrahams V, Guller S. 2011. Inflammation and pregnancy: the role of the immune system at the implantation site. Ann N Y Acad Sci. 1221:80–87.

Mordecai KL, Rubin LH, Maki PM. 2008. Effects of menstrual cycle phase and oral contraceptive use on verbal memory. Horm Behav. 54:286–293.

Mosconi L, Berti V, Quinn C, McHugh P, Petrongolo G, Varsavsky I, Osorio RS, Pupi A, Vallabhajosula S, Isaacson RS. 2017. Sex differences in Alzheimer risk: Brain imaging of endocrine vs chronologic aging. Neurology. 89:1382–1390.

Mott NN, Pak TR. 2013. Estrogen Signaling and the Aging Brain: Context-Dependent Considerations for Postmenopausal Hormone Therapy. ISRN Endocrinol. 2013:1–16.

Ning K, Zhao L, Franklin M, Matloff W, Batta I, Arzouni N, Sun F, Toga AW. 2020. Parity is associated with cognitive function and brain age in both females and males. Sci Rep. 1–9.

Nir T, Villalon-Reina J, Zhu A, Thompson P, Kochunov P, Jahanshad N. 2021. Sensitivity of NODDI Microstructural Measures to the Effects of Age With and Without White Matter Skeletonization. Biol Psychiatry. 89:S278.

Nir TM, Jahanshad N, Villalon-Reina JE, Isaev D, Zavaliangos-Petropulu A, Zhan L, Leow AD, Jack Jr CR, Weiner MW, Thompson PM, (ADNI) ADNI. 2017. Fractional anisotropy derived from the diffusion tensor distribution function boosts power to detect Alzheimer’s disease deficits. Magn Reson Med. 78:2322–2333.

Nir TM, Jahanshad N, Villalon-Reina JE, Isaev D, Zavaliangos-Petropulu A, Zhan L, Leow AD, Jack CR, Weiner MW, Thompson PM. 2017. Fractional anisotropy derived from the diffusion tensor distribution function boosts power to detect Alzheimer’s disease deficits. Magn Reson Med. 78:2322–2333.

Nobis L, Manohar SG, Smith SM, Alfaro-Almagro F, Jenkinson M, Mackay CE, Husain M. 2019. Hippocampal volume across age: Nomograms derived from over 19,700 people in UK Biobank. Neuroimage Clin. 23:101904.

Pierpaoli C, Jezzard P, Basser PJ, Barnett A, Di Chiro G. 1996. Diffusion tensor MR imaging of the human brain. Radiology. 201:637–648.

Pievani M, Filippini N, van den Heuvel MP, Cappa SF, Frisoni GB. 2014. Brain connectivity in neurodegenerative diseases—from phenotype to proteinopathy. Nat Rev Neurol. 10:620–633.

Pines AR, Cieslak M, Larsen B, Baum GL, Cook PA, Adebimpe A, Dávila DG, Elliott MA, Jirsaraie R, Murtha K, Oathes DJ, Piiwaa K, Rosen AFG, Rush S, Shinohara RT, Bassett DS, Roalf DR, Satterthwaite TD. 2020. Leveraging multi-shell diffusion for studies of brain development in youth and young adulthood. Dev Cogn Neurosci. 43.

Pletzer BA, Kerschbaum HH. 2014. 50 Years of Hormonal Contraception - Time To Find Out, What It Does To Our Brain. Front Neurosci. 8:1–6.

Reeves S, Brown R, Howard R, Grasby P. 2009. Increased striatal dopamine (D2/D3) receptor availability and delusions in Alzheimer disease. Neurology. 72:528–534.

Resnick SM, Pham DL, Kraut MA, Zonderman AB, Davatzikos C. 2003. Longitudinal magnetic resonance imaging studies of older adults: a shrinking brain. J Neurosci. 23:3295–3301.

Ritchie SJ, Cox SR, Shen X, Lombardo M v, Reus LM, Alloza C, Harris MA, Alderson HL, Hunter S, Neilson E, Liewald DCM, Auyeung B, Whalley HC, Lawrie SM, Gale CR, Bastin ME, Mcintosh AM, Deary IJ. 2018. Sex Differences in the Adult Human Brainlll: Evidence from 5216 UK Biobank Participants. 2959–2975.

Royston P, Altman DG. 1994. Regression Using Fractional Polynomials of Continuous Covariates: Parsimonious Parametric Modelling. Appl Stat. 43:429.

Salminen LE, Tubi MA, Bright J, Thomopoulos SI, Wieand A, Thompson PM. 2022. Sex is a defining feature of neuroimaging phenotypes in major brain disorders. Hum Brain Mapp. 43:500–542.

Sauerbrei W, Meier-Hirmer C, Benner A, Royston P. 2006. Multivariable regression model building by using fractional polynomials: Description of SAS, STATA and R programs. Comput Stat Data Anal. 50:3464–3485.

Savolainen-Peltonen H, Rahkola-Soisalo P, Hoti F, Vattulainen P, Gissler M, Ylikorkala O, Mikkola TS. 2019. Use of postmenopausal hormone therapy and risk of Alzheimer’s disease in Finland: nationwide case-control study. bmj.364.

Scicali R, Rosenbaum D, Di Pino A, Giral P, Cluzel P, Redheuil A, Piro S, Rabuazzo AM, Purrello F, Bruckert E. 2018. An increased waist-to-hip ratio is a key determinant of atherosclerotic burden in overweight subjects. Acta Diabetol. 55:741–749.

Shumaker SA, Legault C, Rapp SR, Thal L, Wallace RB, Ockene JK, Hendrix SL, Jones III BN, Assaf AR, Jackson RD, Morley Kotchen J, Wassertheil-Smoller S, Wactawski-Wende J, for the WHIMS Investigators. 2003. Estrogen Plus Progestin and the Incidence of Dementia and Mild Cognitive Impairment in Postmenopausal Women. JAMA. 289:2651.

Simerly RB. 2002. Wired for reproduction: organization and development of sexually dimorphic circuits in the mammalian forebrain. Annu Rev Neurosci. 25:507–536.

Smith GD, Whitley E, Dorling D, Gunnell D. 2001. Area based measures of social and economic circumstances: cause specific mortality patterns depend on the choice of index. J Epidemiol Community Health (1978). 55:149–150.

Smith SM, Jenkinson M, Johansen-Berg H, Rueckert D, Nichols TE, Mackay CE, Watkins KE, Ciccarelli O, Cader MZ, Matthews PM. 2006. Tract-based spatial statistics: voxelwise analysis of multi-subject diffusion data. Neuroimage. 31:1487–1505.

Song Y, Li S, Li X, Chen X, Wei Z, Liu Q, Cheng Y. 2020. The Effect of Estrogen Replacement Therapy on Alzheimer’s Disease and Parkinson’s Disease in Postmenopausal Women: A Meta-Analysis. Front Neurosci. 14:1–13.

Spillantini MG, Goedert M. 2013. Tau pathology and neurodegeneration. Lancet Neurol. 12:609–622.

Thomopoulos SI, Nir TM, Reina JEV, Jahanshad N, Thompson PM. 2020. Diffusion MRI metrics of brain microstructure in Alzheimer’s disease: Boosting disease sensitivity with multi-shell imaging and advanced pre-processing. Alzheimer’s & Dementia. 16:1–4.

Thomopoulos SI, Nir TM, Villalon-reina JE, Zavaliangos-petropulu A, Maiti P, Zheng H, Nourollahimoghadam E, Jahanshad N, Thompson PM, for the Alzheimer’s Disease Neuroimaging Initiative S 2021. 2021. Diffusion MRI Metrics and their Relation to Dementia Severitylll: Effects of Harmonization Approaches. SIPAIM. submitted.

Toschi N, Gisbert RA, Passamonti L, Canals S, de Santis S. 2020. Multishell diffusion imaging reveals sex-specific trajectories of early white matter degeneration in normal aging. Neurobiol Aging. 86:191–200.

Townsend JD, Bookheimer SY, Foland-Ross LC, Moody TD, Eisenberger NI, Fischer JS, Cohen MS, Sugar CA, Altshuler LL. 2012. Deficits in inferior frontal cortex activation in euthymic bipolar disorder patients during a response inhibition task. Bipolar Disord. 14:442–450.

Veraart J, Novikov DS, Christiaens D, Ades-Aron B, Sijbers J, Fieremans E. 2016. Denoising of diffusion MRI using random matrix theory. Neuroimage. 142:394–406.

Villalon-Reina JE, Ching CRK, Kothapalli D, Sun D, Nir T, Lin A, Forsyth JK, Kushan L, Vajdi A, Jalbrzikowski M. 2018. Alternative diffusion anisotropy measures for the investigation of white matter alterations in 22q11. 2 deletion syndrome. In: 14th International Symposium on Medical Information Processing and Analysis. International Society for Optics and Photonics. p. 109750U.

World Health Organization. 2019. WHO global report on traditional and complementary medicine 2019. World Health Organization.

Zhang H, Schneider T, Wheeler-Kingshott CA, Alexander DC. 2012. NODDI: Practical in vivo neurite orientation dispersion and density imaging of the human brain. Neuroimage. 61:1000–1016.

