## Supplemental Material for "Exogenous Sex Hormone Effects on Brain Microstructure in Women: A diffusion MRI Study in the UK Biobank"

**Diffusion tensor imaging (DTI)**

Diffusion tensor imaging (DTI) (Basser et al. 1994) fitting was performed on the single-shell (b=1000 s/mm2, 50 diffusion-encoding directions) diffusion MRI data using the DTI fitting (DTIFIT) tool via FSL (<http://fsl.fmrib.ox.ac.uk/fsl/fsl4.0/fdt/fdt_dtifit.html>). A single diffusion tensor, a 3D ellipsoid, capturing a single fiber orientation was modeled at each voxel of the brain. This model assumes a 3D gaussian diffusion process fitted using six independent components, 3 eigenvalues and 3 eigenvectors, describing shape and orientation of the tensor model, respectively. The DTI approach was used to create maps of tensor-derived fractional anisotropy (FA^DTI^) (Jones 2008), representing the fraction of the tensor that can be assigned to anisotropic diffusion and is often considered to measure the degree of directional coherence within a fiber bundle:

$$FA=\sqrt{\frac{3}{2}}\frac{\sqrt{(\lambda_{1}- <\lambda>)^{2}+ (\lambda_{2}- <\lambda>)^{2}+ (\lambda_{3}- <\lambda>)^{2}}}{\sqrt{\lambda_{1}^{2}+\lambda_{2}^{2}+\lambda_{3}^{2}}}\epsilon[0,1]$$

The magnitude of the tensor is defined by the sum of the squares of its eigenvalues. The average diffusivity in all directions, or mean diffusivity (MD^DTI^), is represented by < and >. The primary (largest) eigenvalue, measuring the diffusivity parallel to the majority of axonal fibers, defines axial diffusivity (AD^DTI^). The average between the second and third eigenvalues (λ2 and λ3) defines radial diffusivity (RD^DTI^).The FA^DTI^ index is normalized to take values from 0, when diffusion is isotropic, or 1, when constrained along one axis. In this study, we also considered MD^DTI^, AD^DTI^ and RD^DTI^ maps in addition to FA^DTI^ maps.

**Tensor distribution function (TDF)**

Single-shell diffusion MRI data (b=1000 s/mm, 50 diffusion-encoding directions) was used as input data to estimate the tensor distribution function (TDF) at each voxel in the image (Leow et al. 2009; Nir, Jahanshad, Villalon-Reina, Isaev, Zavaliangos-Petropulu, Zhan, Leow, Jack, et al. 2017). Unlike other models, TDF does not require the specification of the total number of compartments in a tissue, rather it relies on a continuous distribution of tensors, together with a distribution of weights or probabilities. The TDF approach describes intravoxel fibers mathematically as a probabilistic collection of tensors, the probability distribution function P(D), which is defined on a continuous mixture of Gaussian densities in tensor space ($\mathbb{D}$). Here 𝑆(𝑞) represents the measured image intensity signal via the *Fourier transform*:

$$S\left( q \right)=\int_{D\in\mathbb{D}} P(D)e^{(-{tq}^{T}D{d)}_{dD}}$$

Here the wave-vector 𝑞 = 𝑟𝛿𝐺, where r, 𝛿 and 𝐺 being a function of the gyromagnetic ratio, the duration of the diffusion sensitization, and the applied magnetic gradient vector, respectively.

The local maxima of the tensor orientation distribution (TOD) is examined to estimate the number of detected peaks, and defined in the unit sphere along directions θ, TOD(θ):

$$TOD\left( \theta\right)=p(D\left( \lambda,\theta\right))d\lambda$$

For each θ, TDF eigenvalues (λ) of each fiber are calculated by computing the expected values along the principal direction of the fiber. Follows the computation of a scalar TDF anisotropy measure, FA^TDF^, calculated as an extension of the standard FA formula:

$$TDF-FA=TOD\left( \theta\right)*FA(\theta)d\theta$$

$$=\frac{\sqrt{({\lambda^{'}}_{1}\left( \theta\right)-{\lambda^{'}}_{2}\left( \theta\right))^{2}+ ({\lambda^{'}}_{1}\left( \theta\right)-{\lambda^{'}}_{3}\left( \theta\right))^{2}+({\lambda^{'}}_{2}\left( \theta\right)-{\lambda^{'}}_{3}\left( \theta\right))^{2}}}{2[{\lambda^{'}}_{1}\left( \theta\right)^{2}+{\lambda^{'}}_{2}\left( \theta\right)^{2}{{+\lambda}^{'}}_{3}\left( \theta\right)^{2}]}$$

$$where {\lambda^{'}}_{1}\left( \theta\right)=\frac{\int P(D\left( \theta, \lambda\right))\lambda_{i}d\lambda}{\int P(D\left( \theta, \lambda\right))d\lambda}$$

The FA^TDF^ measure differs from the standard FA^DTI^ measure, as it is aimed at adjusting for partial volume effects and weight contribution in regions of fibers mixing or crossing where traditional tensor models tend to fail.

**Neurite Orientation Dispersion and Density Imaging (NODDI)**

In addition to DTI and TDF models, multi-shell dMRI data (b=1000 and 2000 s/mm, 100 diffusion-encoding directions total) were input into Neurite Orientation Dispersion and Density Imaging (NODDI) model, using the AMICO (Accelerated Microstructure Imaging via Convex Optimization) tool. NODDI models the dMRI signal *S*(q) as consisting of intra-cellular, extra-cellular, and isotropic water components; each of these components uniquely affects diffusion in the environment and results in a separate dMRI signal (Zhang et al. 2012).

$$S(q)= v_{iso}S_{iso}\left( q \right)+\left( 1-v_{iso} \right)\left[ v_{ic}S_{ic}\left( q \right)+v_{ec}S_{ec}\left( q \right) \right]$$

The dMRI signal of the intra-cellular, extra-cellular, and isotropic compartments are represented here as S_ic_, S_ec_, S_iso_, respectively. The corresponding volume fractions are v_ic_, v_ec_, v_iso_, where v_ec_ = 1 – _vic_ . S_ic_ is modeled as a set of dispersed cylinders of zero radius, where the dispersion is defined by a Watson distribution. S_ec_ represents a dispersed mixture of Gaussian anisotropic diffusion, and E_iso_ represents isotropic Gaussian diffusion (Zhang et al. 2012; Daducci et al. 2015). We examined the following voxel-wise microstructural parameters: ODI^NODDI^ (orientation dispersion index, a measure of within-voxel tract white matter disorganization), ICVF^NODDI^ (intracellular volume fraction, and index of white matter neurite density) and ISOVF^NODDI^ (isotropic or free water volume fraction) (Zhang et al. 2012; Daducci et al. 2015).

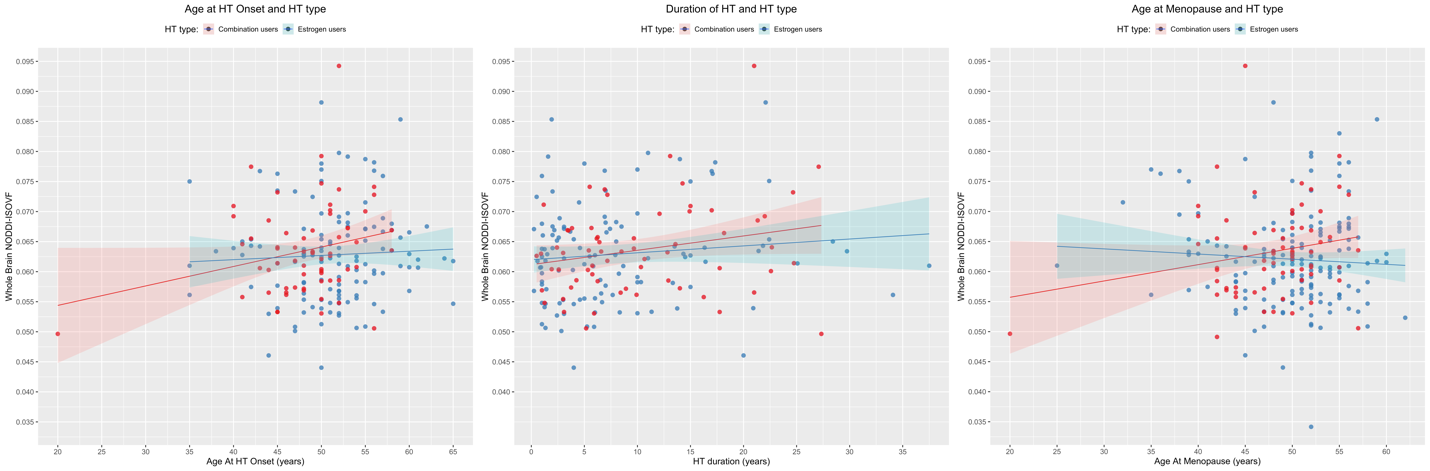

**Figure 1.**  **Age at HT onset, duration of HT, and age at menopause for ISOVF^NODDI^ in estrogen only users and combination users.** In combination HT (estrogen + progestin) users, later age at HT onset (*left panel*), prolonged duration of HT therapy (*middle panel*), and later age at menopause (*right panel*), were associated with higher ISOVF^NODDI^, relative to estrogen only HT users. Blue colors indicate estrogen only HT users, red colors indicate combination HT users. Dotted lines represent 95% confidence intervals. Of note, subjects with younger age at HT onset, and age at menopause had either undergone surgeries or had history of cancer.

Basser, P. J., Mattiello, J., & LeBihan, D. (1994). MR diffusion tensor spectroscopy and imaging. *Biophysical Journal*, *66*(1), 259–267. https://doi.org/10.1016/S0006-3495(94)80775-1

Daducci, A., Canales-Rodríguez, E. J., Zhang, H., Dyrby, T. B., Alexander, D. C., & Thiran, J.-P. (2015). Accelerated microstructure imaging via convex optimization (AMICO) from diffusion MRI data. *NeuroImage*, *105*, 32–44.

Leow, A. D., Zhu, S., Zhan, L., McMahon, K., de Zubicaray, G. I., Meredith, M., Wright, M. J., Toga, A. W., & Thompson, P. M. (2009). The tensor distribution function. *Magnetic Resonance in Medicine*, *61*(1), 205–214. https://doi.org/10.1002/mrm.21852

Nir, T. M., Jahanshad, N., Villalon-Reina, J. E., Isaev, D., Zavaliangos-Petropulu, A., Zhan, L., Leow, A. D., Jack, C. R., Weiner, M. W., & Thompson, P. M. (2017). Fractional anisotropy derived from the diffusion tensor distribution function boosts power to detect Alzheimer’s disease deficits. *Magnetic Resonance in Medicine*, *78*(6), 2322–2333. https://doi.org/10.1002/mrm.26623

Zhang, H., Schneider, T., Wheeler-Kingshott, C. A., & Alexander, D. C. (2012). NODDI: practical in vivo neurite orientation dispersion and density imaging of the  human brain. *NeuroImage*, *61*(4), 1000–1016. https://doi.org/10.1016/j.neuroimage.2012.03.072
